# Targeting pediatric High-Grade Gliomas with *O*AcGD2-CAR Vδ2 T cells

**DOI:** 10.1101/2023.11.17.567375

**Authors:** Pauline Thomas, Maëva Veerasamy, Marine Devinat, Elodie Guiet, Jocelyn Ollier, Pierre Paris, Natacha Entz-Werlé, Catherine Gratas, Béatrice Clémenceau, Stéphane Birklé, François Paris, Claire Pecqueur, Sophie Fougeray

**Author notes:** Corresponding author: Sophie Fougeray CRCI2NA, Nantes Université, INSERM U1307, CNRS 6075, 8 quai Moncousu 44007 Nantes, France. **E-mail:**. **Translational relevance** Pediatric high-grade gliomas (pHGG) are a family of rare cancers of childhood with poor prognosis despite medical advances. T cells expressing chimeric antigen receptor (CAR) are nowadays evaluated to efficiently fight these tumors. Nevertheless, all studies are investigating conventional αβ T cells autologous transfer targeting tumor antigens that are usually also expressed in normal tissues. Here, we demonstrated that OAcGD2, an antigen that is restricted to tumors, is expressed in pHGG and can be targeted by *O*AcGD2-CAR T cells. In addition to conventional αβ OAcGD2 CAR-T cells, we also propose a strategy using Vδ2-CAR T cells, which exhibited a similar anti-tumor reactivity compared to αβ T cells without any allogeneic reactivity allowing banking and off-the-shelf CAR-T cell therapy.

## Abstract

**Purpose:** Pediatric high-grade gliomas (pHGG) belong to a family of rare children’s cancers which are treated with radiotherapy, based on adult high-grade glioma standard of care. However, new treatments are definitively required since actual ones are unable to extend survival by more than a few months in most patients. In this study, we investigate a Chimeric Antigen Receptor (CAR)-T cell immunotherapy targeting the *O*AcGD2 ganglioside, using either conventional αβ or Vδ2 T cells as effectors.

**Materials and methods:** Using relevant human primary models of pHGG, we first characterized the innate Vδ2 T cell immunoreactivity. Then, following the validation of *O*AcGD2 expression in these tumor cells, we evaluated both αβ and Vδ2 *O*AcGD2-CAR-T cell immunoreactivity using various methods including videomicroscopy, FACS and cytotoxicity assays.

**Results:** We showed that pHGG primary cells are not spontaneously recognized and killed by Vδ2 T cells but significantly expressed the *O*AcGD2 ganglioside. Accordingly, both αβ and Vδ2 T cells engineered to express a CAR against the *O*AcGD2 efficiently killed pHGG cells in 2D and 3D models. Importantly, only Vδ2 T cells transduced with the complete *O*AcGD2-CAR eliminated pHGG cells, in contrast to conventional αβ CAR-T cells that killed tumor cells even in the absence of CAR expression, highlighting the allogeneic potential of Vδ2 CAR-T cells.

**Conclusion:** Our study demonstrates the preclinical relevance of targeting *O*AcGD2 in pHGG using CAR-T cells. Furthermore, we also clearly demonstrate the clinical benefits of using Vδ2 T cells as CAR effectors in allogeneic settings allowing an off-the-shelf immunotherapy.

## Introduction

Pediatric brain tumors, including pediatric high-grade gliomas (pHGG), are the leading cause of cancer-related death in childhood. Pediatric HGG include diffuse hemispheric gliomas (DHG) and diffuse midline gliomas (DMG). The actual prognosis for pHGG patients remains drastically poor since current therapeutic strategies are unable to extend survival by more than a few months in most patients. Notably, diffuse intrinsic pontine glioma (DIPG), belonging to DMG, is still the only child’s cancer with no curative treatment with a median survival of less than one year (1). Current treatments are based on clinical protocols for adult high-grade glioma (aHGG) and mostly involve a combination of radio- and chemotherapies. However, the WHO classification in 2021 clearly sets apart pHGG from aHGG (2). The molecular profiling of these tumors also led to the development of immunotherapy as an emerging treatment for pHGG (NCT04185038, NCT03500991, NCT03638167, NCT05298995). While some promising results have been observed in some clinical studies (3,4), all targeted antigens are also expressed in normal tissues and may result in *on-target/off-tumor* effects. Interestingly, we have identified *O*AcGD2, the *O-*acetylated form of the GD2 ganglioside, as a specific tumor antigen in various tumors of neuro-ectodermic origin (5–8). Furthermore, its targeting using a monoclonal antibody reduced tumor progression in neuroblastoma and aHGG preclinical models (6,9,10).

In solid cancers, immunotherapy usually involves conventional αβ CAR-T cells. However, manufacturing of such effectors may rapidly become challenging in patients treated with radiochemotherapy. In contrast, T cells from a healthy donor may provide a good source of immune cells for adoptive immunotherapy and can be used to generate off-the-shelf CAR T cells that are readily available for administration into patients when required. Interestingly, Vδ2 T cells do not recognize MHC-I molecules allowing an allogeneic administration. They rather get activated upon encounter with transformed cells or stressed cells, for example following radiotherapy (11). Moreover, we recently revealed the innate recognition of Vδ2 T cells toward a subset of aHGG cells (12). Indeed, aHGG expressing significant levels of butyrophilins and NKG2D ligands, which are the respective ligands of Vδ2 TCR and NKG2D receptors, are efficiently killed by Vδ2 T cells both *in vitro* and in an orthotopic model of aHGG.

In this study, we evaluated the antitumor potential of *O*AcGD2-CAR T cells against pHGG. We demonstrated that, while we detected a weak innate recognition of Vδ2 T cells, all pHGG cells, including DIPG, expressed significant expression of *O*AcGD2. Using both 2D and 3D tumoroid models, we further demonstrated that *O*AcGD2-CAR engineering allowed tumor cell killing using both conventional αβ T cells and Vδ2 T cells. Finally, we underlined the allogeneic potential of *O*AcGD2-CAR Vδ2 T cells in our 3D models. Altogether, our study paved the way for developing allogeneic immunotherapy using *O*AcGD2-CAR Vδ2 T cells to treat pHGG.

## Materials and Methods

The patient-derived primary cultures, and the human CHLA-01-MED medulloblastoma cell line purchased from ATCC, were cultured in neurobasal media, as previously described (13). The human DHG cell line was purchased from the Children’s Oncology Group and cultured in complete media. T cells were expanded from PBMCs obtained from seven blood donors at the Etablissement Franc□ais du Sang (EFS) with informed consent (Blood product transfer agreement relating to biomedical research protocol 97/5-B—DAF 03/4868). αβ and Vδ2 T cells were respectively expanded with CD3/CD28 beads and Zoledronate, both in presence of IL-2. T cells were retrovirally transduced to express the CAR with (ζCAR) or without (ΔCAR) the CD3ζ transducing domain (6). RNA expression was evaluated by quantitative real-time PCR (qPCR) using an aHGG primary culture as a control. Cell phenotypes and T cell degranulation were measured by flow cytometry (Accuri C6Plus, Canto II, Symphony, BD). Cytolytic activity was performed by ^51^Cr assay in 2D models and by videomicroscopy in 3D models using an Incucyte® (Sartorius, Germany). All results are presented as mean ± SD from independent experiments, as described in the corresponding legends. Statistical analyses were performed using Prism 9.0 GraphPad Software. For detailed methods, see STAR methods file.

## Results

### Vδ2 T cells displayed a weak innate reactivity against pHGG tumor cells

To evaluate Vδ2 T cell abilities to naturally recognize pHGG, we measured the relative RNA expression of the Vδ2 TCR ligands (BTN2A1 and BTN3A1) and NKG2D-ligands (ULBP2, MICA/B) in one DHG cell line and two DIPG primary cultures (**Figure 1A**). An aHGG primary culture known to be spontaneously recognized by Vδ2 T cells was used as a control. All primary cultures displayed heterogeneous expression of Vδ2 TCR and NKG2D-ligands. In general, all ligands were less expressed in pHGG cells as compared with aHGG cells. Furthermore, while pHGG expressed significant levels of BTN2A1 and BTN3A1, they barely expressed NKG2D-ligands. Similar results were obtained at the protein levels (**Figure 1B**, 0 Gy). Since radiotherapy is known to induce NKG2D-ligand expression, their expression was assessed 72 hours following increasing doses of irradiation (**Figure 1B**). Surprisingly, neither ULBP2 nor MICA/B expression was altered by irradiation. Expression of other inhibiting (HLA-E) and activating (PVR) ligands was also measured after irradiation (**Figure 1B**). HLA-E expression tended to increase with irradiation doses while a significant decrease in PVR expression was observed after 10 and 15 Gy irradiation in the 3 pHGG cell lines. To determine how these observations translate into antitumor functions of Vδ2 T cells, we measured Vδ2 T cell activation by FACS analysis through the expression of the degranulation marker CD107a following their co-culture with tumor cells (**Figure 1C and D**). In agreement with the low expression of Vδ2 TCR and NKG2D-ligands, Vδ2 activation was weak in the presence of pHGG cells. Surprisingly, activation of Vδ2 T cells was stronger with DIPG-1 primary cells, as compared with DHG and DIPG-2 cells. As expected, Vδ2 T cells got activated in the presence of aHGG cells. Irradiation did not affect Vδ2 T cell activation (**Figure 1D**). Altogether, these results show that Vδ2 T cells displayed a weak innate reactivity against pHGG cells.

**Figure 1:**
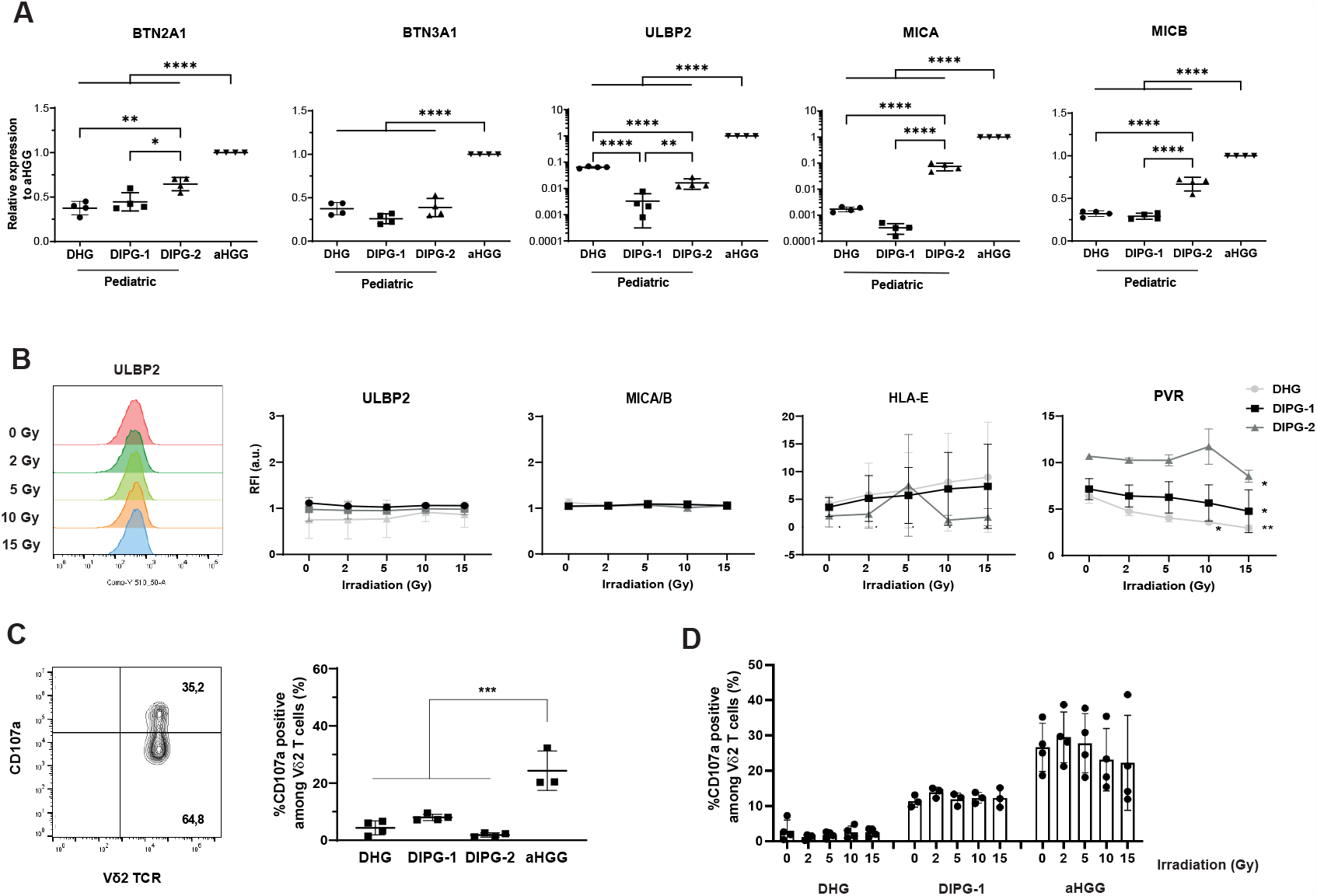
Vδ2 T cells slightly recognize pHGG cells through innate reactivity. **A**. Expression levels of BTN2A1, BTN3A1, MICA, MICB and ULBP2 mRNA in DHG, DIPG-1 and DIPG-2 tumor cells, measured by RT-qPCR, in comparison with mesenchymal adult High-Grade Glioma (aHGG) expression. Results are expressed as mean ± SD, n=4. Two-way ANOVA test, Tukey’s multiple comparisons test *, p<0.05 ; **, p<0.01 ; ****, p<0.0001. **B**. Expression of several Vδ2 TCR ligands following irradiation. Representative flow cytometry histogram of ULBP2 expression in DHG cells 72 hours after 0, 2, 5, 10 and 15 Gy (left panel). Ratio of Fluorescence Intensity (RFI) of ULBP2, MICA/B, HLA-E and PVR expression in DHG cell line, DIPG-1 and DIPG-2 72 hours after 0, 2, 5, 10 or 15 Gy irradiation (right panel). Results are presented as mean ± SD, n=3. Two-way ANOVA test, Dunnett’s multiple comparisons test, *, p<0.05 ; **, p<0.01. **C**. Activation of Vδ2 T cells. Representative histogram (left panel) and frequency of CD107a+ cells among Vδ2+ T cells (right panel). Vδ2 T cell activation was tested with V δ2 T cells from different donors cocultured with DHG cell line, DIPG-1, DIPG-2 primary cells or aHGG primary cells at an effector:target ratio 1:1. Results are presented as mean ± SD, n ≥ 3 donors; for each donor n ≥ 2. One-way ANOVA test, Tukey’s multiple comparison test, ***, p<0.001. **D**. Frequency of CD107a+ cells among Vδ2+ T cells from different donors cocultured with DHG cell line, DIPG-1 primary cells or aHGG primary cells at 1:1 effector:target ratio, 72 hours after tumor cell irradiation (0, 2, 5, 10 or 15 Gy). Results are presented as mean ± SD, n ≥ 3 donors ; for each donor n ≥ 2.

### *O*AcGD2 ganglioside is significantly expressed on pHGG

To establish the relevance of *O*AcGD2 targeting, its expression was determined in DHG and DIPG primary cells (**Figure 2A and B**). Similar levels of *O*AcGD2 were observed in all 3 primary cultures. Similar results were observed in two other DIPG and one DHG primary cells, BT68NS, BT69NS, and BT35 respectively (**Figure 2B, Extended Table 1**). *O*AcGD2 expression was also determined 72 hours following increasing doses of irradiation. Importantly, irradiation did not affect *O*AcGD2 expression on DIPG primary cells (**Figure 2C**). Thus, pHGG cells express significant levels of *O*AcGD2, which are not affected by irradiation.

**Figure 2:**
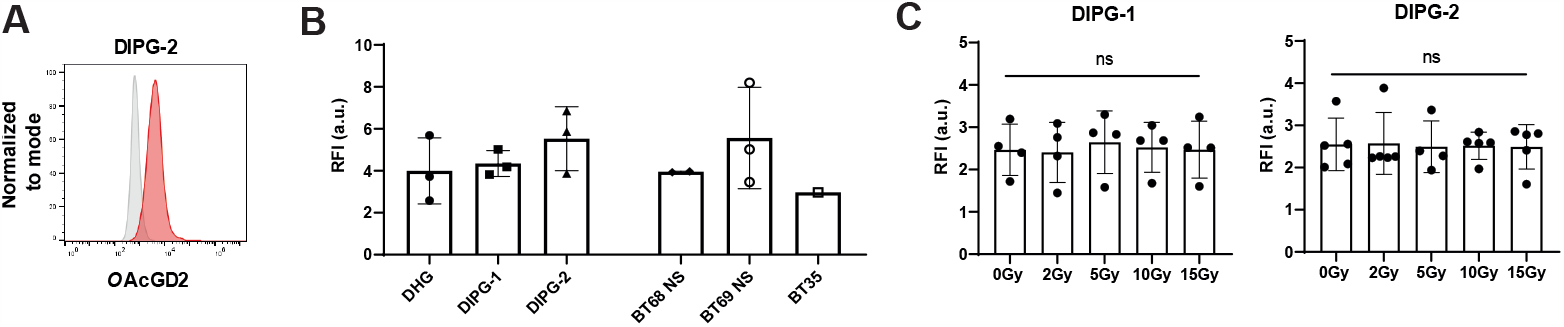
*O*AcGD2 ganglioside is expressed on pediatric High-Grade Glioma cells. **A**. Representative flow cytometry histogram of *O*AcGD2 expression on DIPG-2 primary culture. **B**. Ratio of Fluorescence Intensity (RFI) of *O*AcGD2 expression on pHGG cells. RFI corresponds to the ratio of anti-*O*AcGD2 mAb fluorescence normalized to isotype fluorescence. Results are presented as mean ± SD, n = 3. **C**. Ratio of Fluorescence Intensity (RFI) of *O*AcGD2 expression in DIPG-1 and DIPG-2 72 hours after irradiation (0, 2, 5, 10 or 15 Gy). RFI is calculated by ratio of anti-*O*AcGD2 mAb fluorescence normalized to isotype fluorescence. Results are presented as mean ± SD, n = 4. Two-way ANOVA test.

### *O*AcGD2-CAR-T cells efficiently kill pHGG

We then designed a CAR containing a human mutated IgG1 as the spacer, CD28 as the costimulatory domain, and CD3ζ as the transducing domain (**Figure 3A**). Two different constructions were produced, a complete and effective ζ*O*AcGD2-CAR, and a Δ*O*AcGD2-CAR, lacking the transducing domain, as a negative control. Both αβ and Vδ2 T cells were selectively transduced, sorted, and expanded. More than 85% of αβ T cells were transduced with either Δ*O*AcGD2 and ζ*O*AcGD2-CAR (**Figure 3B**). CAR transduction was slightly less efficient in Vδ2 T cells but still reached 70% of efficacy. To determine whether CAR expression impacted T cell phenotype, we evaluated the expression of several checkpoint inhibitors before and after CAR transduction (**Extended Figure 1A, 1B**). No significant change was observed in the exhaustion profile of αβ or Vδ2 T CAR-T, either in the number of markers expressed by the cells or the percentage of cells expressing Lag3, PD1, TIGIT, or Tim-3. We further assessed CAR-T cell phenotype through the expression of either CD8/CD4 on αβ T cells or NKG2D/DNAM-1/NKG2A on Vδ2 T cells. αβ T cells were enriched in CD8+ cells after ζ*O*AcGD2-CAR transduction, as compared to non-transduced or ΔCAR T cells (**Extended Figure 1C**). Transduction of ζ*O*AcGD2-CAR resulted in a decrease in NKG2D receptor and DNAM-1 expression in Vδ2 T cells, while expression of the inhibitory NKG2A receptor was increased (**Extended Figure 1D**). Additionally, both populations displayed an effector memory (Tem) or effector memory re-expressing CD45RA (Temra) (**Extended Figure 1E**). Finally, the ability of CAR-T cells to kill pHGG cells was assessed using degranulation assay and ^51^Cr-release assay (**Figure 3C and D**). As shown by CD107a marker expression, a significant proportion of αβ and Vδ2 T cells got activated in the presence of pHGG cells, when they expressed the effective ζ*O*AcGD2-CAR (**Figure 3C**). In contrast, no activation was observed when T cells were transduced with Δ*O*AcGD2-CAR T cells. Importantly, ζ*O*AcGD2-CAR T cells were not activated when cocultured with CHLA-01-MED cell line that does not express *O*AcGD2 (**Extended Figure 2)**. In agreement with these results, ζ*O*AcGD2-CAR T cells efficiently killed DHG, DIPG-1 and DIPG-2 cells, in a dose-dependent manner (**Figure 3D**). No tumor cell lysis was observed when tumor cells were cocultured with non-transduced (NT) or Δ*O*AcGD2-CAR T cells, or against CHLA-01-MED cells (**Figure 3D**). Despite a significant difference in CD107a expression between αβ and Vδ2 T cells transduced with ζ*O*AcGD2-CAR (17% vs 28%), no difference was observed in tumor cell lysis (**Figure 3E**). Altogether, these results revealed that the transduction of ζ*O*AcGD2-CAR confers to both αβ and Vδ2 T cells the ability to specifically target and kill pHGG tumor cells.

**Figure 3:**
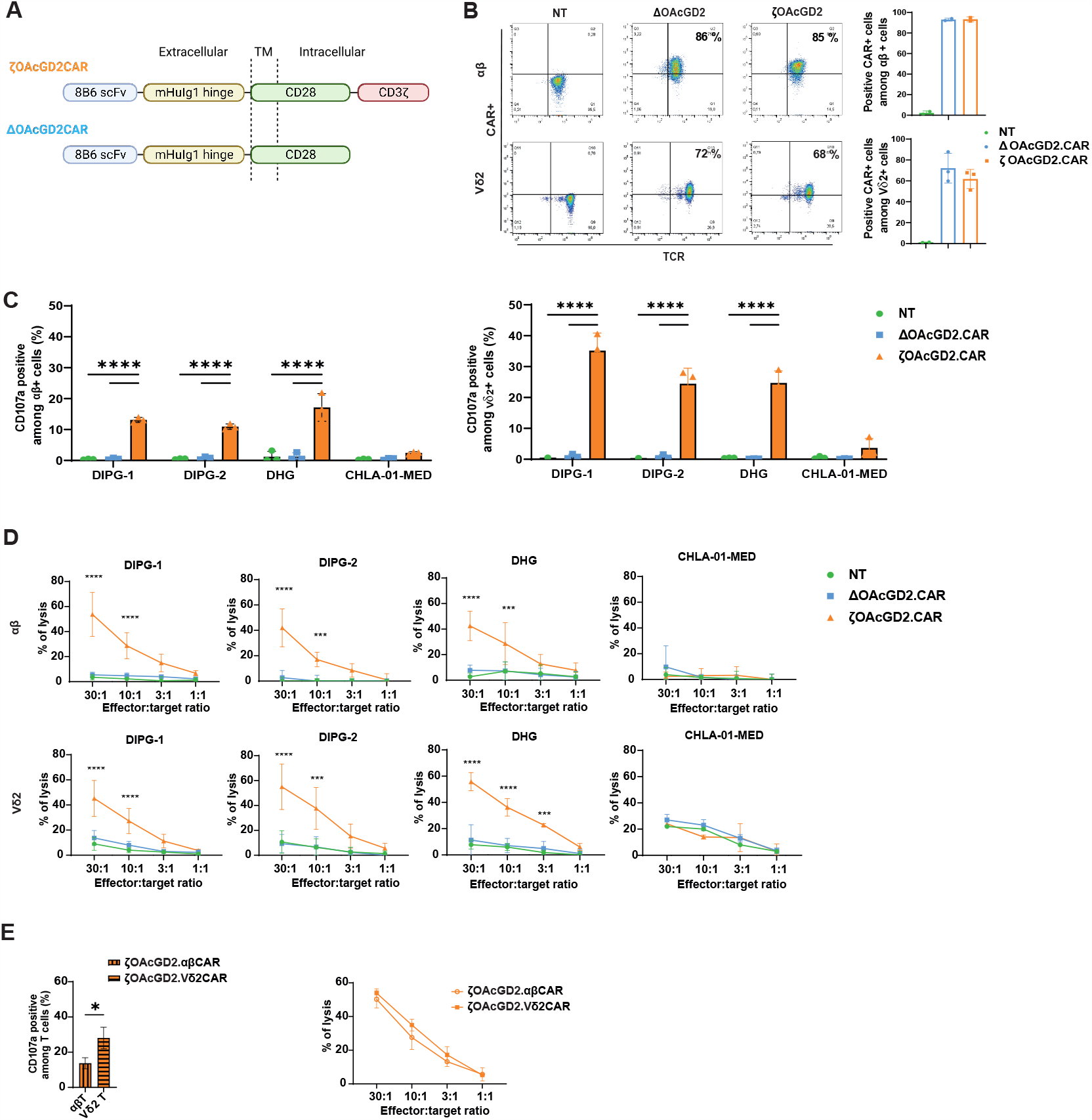
*O*AcGD2-CAR T cells specifically target *O*AcGD2-positive cells. **A**. Generation of *O*AcGD2-CAR. ζ*O*AcGD2-CAR is composed of the anti-*O*AcGD2 mAb-8B6 scFv linked to the mutated human IgG1, the CD28 costimulatory domain and the CD3ζ transducing domain. Δ*O*AcGD2-CAR has the same composition but does not include the transducing domain. **B**. Dot plots (left panel) and expression histograms (right panel) of FACS analysis of transduced-CAR in αβ (top) and Vδ2 (bottom) T cells. Non transduced (NT), Δ*O*AcGD2-CAR and ζ*O*AcGD2-CAR are indicated in green, blue and orange respectively. Results are expressed as % of CAR+ cells within CD3+ αβ or Vδ2+ T cells, n=3. **C**. Frequency of CD107a+ cells among non-transduced, Δ*O*AcGD2-CAR and ζ*O*AcGD2-CAR CD3+ αβ T cells (left panel) or Vδ2+ T cells (right panel) after 4 hours of coculture with 2 DIPG primary cells (DIPG-1 and DIPG-2) and 2 tumor cell lines (DHG and CHLA-01-MED) at an effector:target ratio 1:1. Results are presented as mean ± SD, n=4, two-way ANOVA with Tukey’s multiple comparisons: ****, p<0.0001. **D**. *In vitro* cytotoxicity of non-transduced, Δ*O*AcGD2-CAR and ζ*O*AcGD2-CAR αβ T cells (top) or Vδ2+ T cells (bottom) after 4 hours of coculture with 2 DIPG primary cells (DIPG-1 and DIPG-2) and 2 tumor cell lines (DHG and CHLA-01-MED). Cytotoxicity was evaluated using standard ^51^Cr release assay. Results are expressed as % of cell lysis depending on the indicated effector:target ratio. Results are expressed as mean ± SD, n =4, two-way ANOVA with Tukey’s multiple comparisons: **, p<0.01; ***, p<0.001; ****, p<0.0001. **E**. Frequency of CD107a+ cells (left panel) and tumor cell lysis (right panel) between αβ and Vδ2 CAR-T cells after 4 hours of coculture with pHGG cells (DHG cell line and DIPG-1, DIPG-2 primary cells). Results are presented as mean ± SD, n = 4. T-test: *, p<0.05.

### αβ-CAR-T cells display allogeneic recognition against pHGG, in contrast to Vδ2-CAR-T cells

We further evaluated CAR-T cell efficiency using 3D models of DIPG, using GFP-expressing tumor cells. In these models, *O*AcGD2 expression was mainly detected on the spheroid surface, by confocal microscopy (**Figure 4A**). However, FACS analysis performed just after the dissociation of the tumoroids revealed that *O*AcGD2 was expressed on all tumor cells, without significant differences between 2D and 3D models (**Figure 4B**). Tumor cell survival in 3D models was then monitored with time in the presence of non-transduced, Δ*O*AcGD2-CAR and ζ*O*AcGD2-CAR T cells. All ζ*O*AcGD2-CAR T cells efficiently killed DIPG-2 cells within 2 days, at a ratio of 3:1 effector:target (**Figure 4C and E**). Interestingly, while αβ allogeneic recognition could not be observed in 2D models, both non-transduced and Δ*O*AcGD2-CAR αβ T cells eliminated DIPG-2 tumoroids, in a similar time frame than ζ*O*AcGD2-CAR αβ T cells. In contrast, Vδ2 T cells that were either non-transduced or transduced with the truncated Δ*O*AcGD2-CAR, did not kill DIPG-2 cells (**Figure 4D and E**). Similar results were observed with DIPG-1 cells since all αβ T cells killed DIPG-1 cells whereas only ζ*O*AcGD2-CAR Vδ2 T cells killed them (**Extended Figure 3A, B and C**). Again, no significant changes was observed in the exhaustion profile of αβ and Vδ2 T cells transduced with the ζ*O*AcGD2-CAR, after 5 days of coculture (**Figure 4F and G**). Altogether, our results demonstrate that, while αβ T cells killed pHGG cells in an *O*AcGD2-CAR independent manner, ζ*O*AcGD2-CAR Vδ2 T cells selectively killed pHGG cells in 3D models.

**Figure 4:**
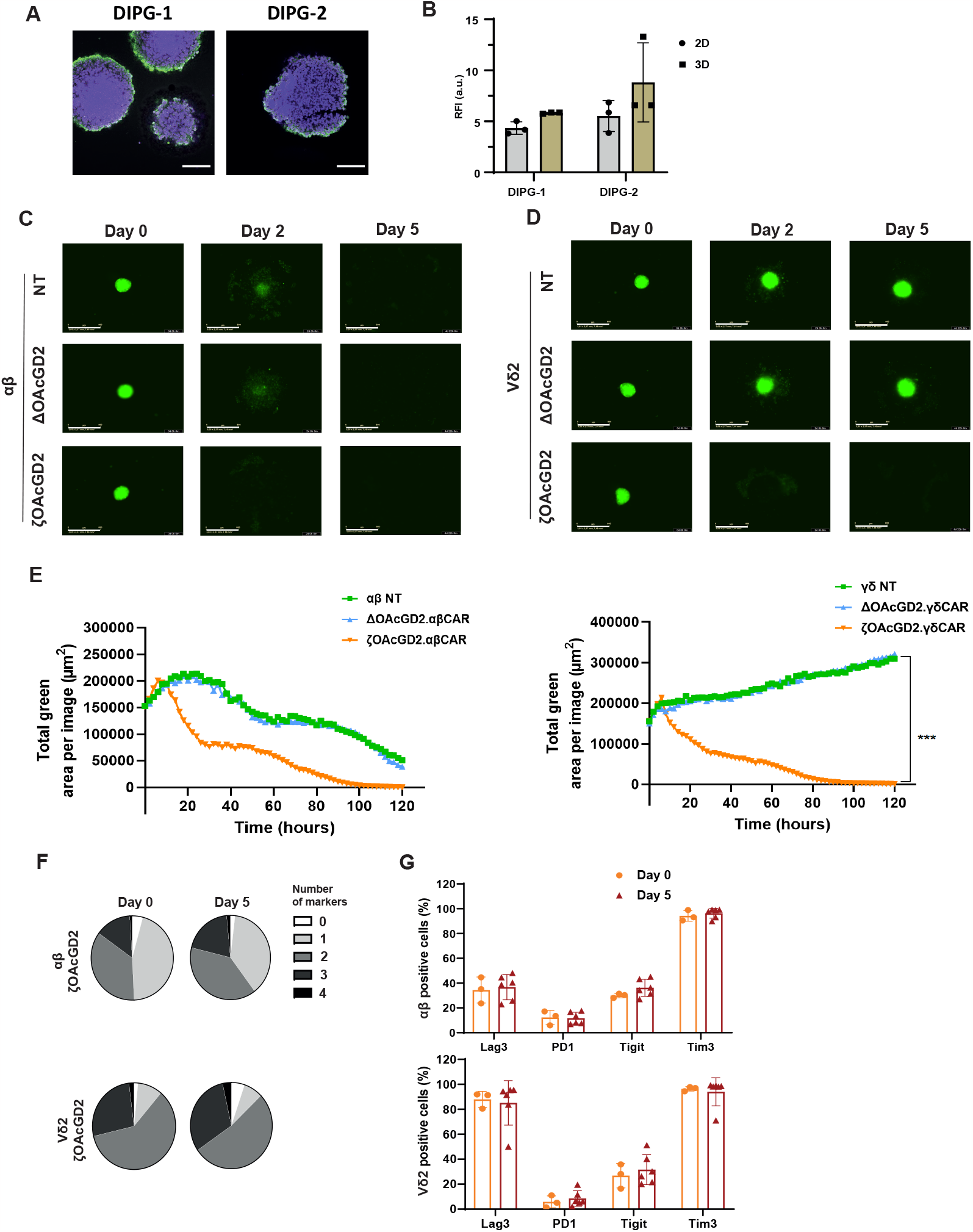
Vδ2 CAR-T cells do not develop long-term allogeneic reactions but induce CAR-cytotoxicity. **A**. Representative picture of *O*AcGD2 expression (green) on DIPG-1 and DIPG-2 in 3D model after transparization using confocal microscopy (nuclei are in violet). Scale bar = 100 μm. **B**. Ratio of Fluorescence Intensity (RFI) of *O*AcGD2 expression on DIPG-1 and DIPG-2 primary cells in single cell suspension from 2D- and 3D-models. Results are presented as mean ± SD, n = 3. **C-D**. Representative images of tumor cell killing over time using GFP-expressing DIPG-2 cells in 3D-model. Pictures were taken on day 0, 2 and 5 following addition of either αβ CAR-T cells (C) or Vδ2 CAR-T cells (D). Immune effectors were non-transduced, transduced with the Δ*O*AcGD2-CAR or the ζ*O*AcGD2 CAR at an effector:target ratio of 3:1. Scale bar = 800 μm. **E**. Representative quantification of tumor cell killing following the addition of immune effectors on 3D models of DIPG-2 at an effector:target ratio of 3:1. Results are presented as mean, n=3, Two-way ANOVA test, ***, p<0.001. **F-G**. Exhaustion profile of αβ (top) and Vδ2 (bottom) CAR-T cells following ζ*O*AcGD2-CAR transduction. Distribution of immune effectors expressing zero, one, two, three or four exhaustion markers (Lag3, PD1, TIGIT and Tim3) (F) and frequency of positive cells expressing exhaustion markers (G) in αβ (top) and Vδ2 (bottom) T cells. Exhaustion profiling was performed at day 0 and 5 of coculture with GFP-expressing DIPG-2 with an effector:target ratio of 3:1. Results are presented as % of positive cells, n = 3.

## Discussion

The remarkable success of CAR-T cell therapies in haematological malignancies have paved the way to develop similar strategies in solid tumors. However, the immune memory induced by CAR-T cells targeting a non-specific tumor antigen may expose patients to *on-target/off-tumor* side effects, which could be life-threatening (14). Prompted by our knowledge demonstrating the specific expression of *O*AcGD2 in tumor cells (6,9), we developed an immunotherapy targeting *O*AcGD2 using CAR T cells against pHGG, incurable pediatric brain cancers. We revealed for the first time the expression of *O*AcGD2 on several pHGG primary cells and demonstrated the relevance of its targeting using *O*AcGD2-CAR-T cells. Importantly, we further underlined the allogeneic potential of *O*AcGD2-CAR Vδ2 T cells since these cells required the expression of the efficient ζ*O*AcGD2-CAR to kill tumor cells, in contrast to conventional αβ T cell effectors that kill them independently of CAR expression. Altogether, our study demonstrates that *O*AcGD2-CAR Vδ2 T cells is a potent strategy that not only specifically targets pHGG tumor cells but also can be used in an allogeneic setting allowing off-the-shelf manufacturing of CAR-T cells.

The *O*AcGD2 ganglioside expression was previously described in aHGG, neuroblastoma and breast cancer (5,6,9). Here, we reported for the first time its expression in pHGG. Importantly, in opposition to GD2, *O*AcGD2 is not expressed in normal tissues, enabling specific tumor cell targeting. Despite the lower level of *O*AcGD2 expression compared to GD2, *O*AcGD2-CAR-T cells were efficiently able to kill tumor cells, in both 2D and 3D models. Moreover, while GD2 scFv has been described to induce CAR-T cell tonic signaling resulting in T cell exhaustion and limited efficiency (15), *O*AcGD2-CAR transduction did not alter T cell exhaustion profile. Altogether, these results favor *O*AcGD2 targeting over GD2 targeting. Expression of *O*AcGD2 remains to be assessed following ζ*O*AcGD2-CAR T cell treatment to determine whether tumor cells might escape CAR-T cell targeting through antigen loss, as commonly observed with protein antigen (16). In this case, the design of a complex CAR targeting several antigens at once might be considered.

Nowadays, the majority of CAR-T cells that have been developed are from conventional αβ T cells, which could be challenging when patients undergo radio-chemotherapeutics treatments, not to mention production time and cost. In our study, we evaluated both αβ and Vδ2 T cell effectors. Both immune effectors displayed similar tumor cell killing abilities and no sign of exhaustion. Similar results were observed by Capsomidis *et al*. which have shown similar *in vitro* efficiency of αβ, Vδ1 and Vδ2 T cells following GD2-CAR transduction (17). However, in our 3D models, we were able to reveal the allogeneic immunoreactivity of αβ T cells that was not observed with Vδ2 T cells, since they do not recognize MHC-I. These results highlighted the great potential of Vδ2 T cells that can be used in allogeneic settings with no risk of Graft versus Host Disease (GvHD), and banked from healthy donors allowing on-demand CAR transduction and “off-the-shelf” CAR therapy (18).

Besides tumor targeting through CAR transduction, Vδ2 T cells display several additional benefits as immune effectors. They displayed a strong and natural cytotoxic potential against several cancers, including aHGG (12,19). Furthermore, conventional treatments such as irradiation or chemotherapy, have been shown to increase NKG2D-ligand expression (11). While Vδ2 T cells display a weak innate immunoreactivity against pHGG cells, even after irradiation, our results do not exclude a clinical synergistic potential effect of Vδ2 TCR and CAR immunoreactivities. Indeed, it would be interesting to determine Vδ2 T cell immunoreactivity against pHGG cells following irradiation protocols that would better mimic the clinical hyper-fractionated protocol proposed to patients. The impact of chemotherapy on Vδ2 T cell immunoreactivity is also worth investigating. Treatment with aminobiphosphonates such as Zoledronate, might also be considered since these pharmacological compounds increase innate Vδ2 T cell immunoreactivity, resulting in better tumor cell killing (20). While Zoledronate has been described as having a poor biodistribution and inducing side-effects when injected systemically (21), several groups are currently investigating innovative formulations. Finally, Vδ2 T cells also display antigen-presenting cells abilities, which further widen their immune functions (17). Yet, Capsomidis *et al*. described how Vδ2 T cells induced the upregulation of CD86 and HLA-DR expression following Zoledronate expansion, increasing their antigen-presenting cell abilities (17).

In conclusion, we have developed *O*AcGD2-CAR-T cells from both conventional αβ and unconventional Vδ2 T cells and demonstrated their clinical relevance against pHGG. Importantly, we provided strong evidence of the clinical potential of Vδ2 CAR-T cells in allogeneic settings and opened a new way of “off-the-shelf” CAR-mediated immunotherapy against DIPG.

## Supporting information

STAR Methods

## Acknowledgements

We thank the Ligue contre le cancer and Fondation ARC for supporting this project. We thank the FACS facility “CytoCell” (SFR François Bonamy), the tissue imaging core facility “Micropicell” (SFR François Bonamy) and the radioactivity technical platform (SFR François Bonamy) for expert technical assistance. Anti-OAcGD2 antibody mAb 8B6 was generously provided by OGD2 Pharma, Nantes, France.

## Authors contributions

SF, CP, SB designed research; NEW provided primary cultures; PT, MV, MD, EG, JO, BC, CG performed research; PT, CG analyzed data; PT, SF and CP wrote the manuscript; all authors revised the manuscript.

## Ethics approval and consent to participate

Informed consent was obtained from all individual participants included in this study and all procedures were in accordance with the ethical standards of the ethic National Research Committee and with the 1964 Helsinki Declaration and its later amendments or comparable ethical standards.

## Data availability statement

All relevant raw data will be freely available to any researcher for non-commercial purposes on request.

