## Extended Figures for "Targeting pediatric High-Grade Gliomas with *O*AcGD2-CAR Vδ2 T cells"

### Extended Figure 1: T cell phenotype is slightly altered by OAcGD2-CAR transduction

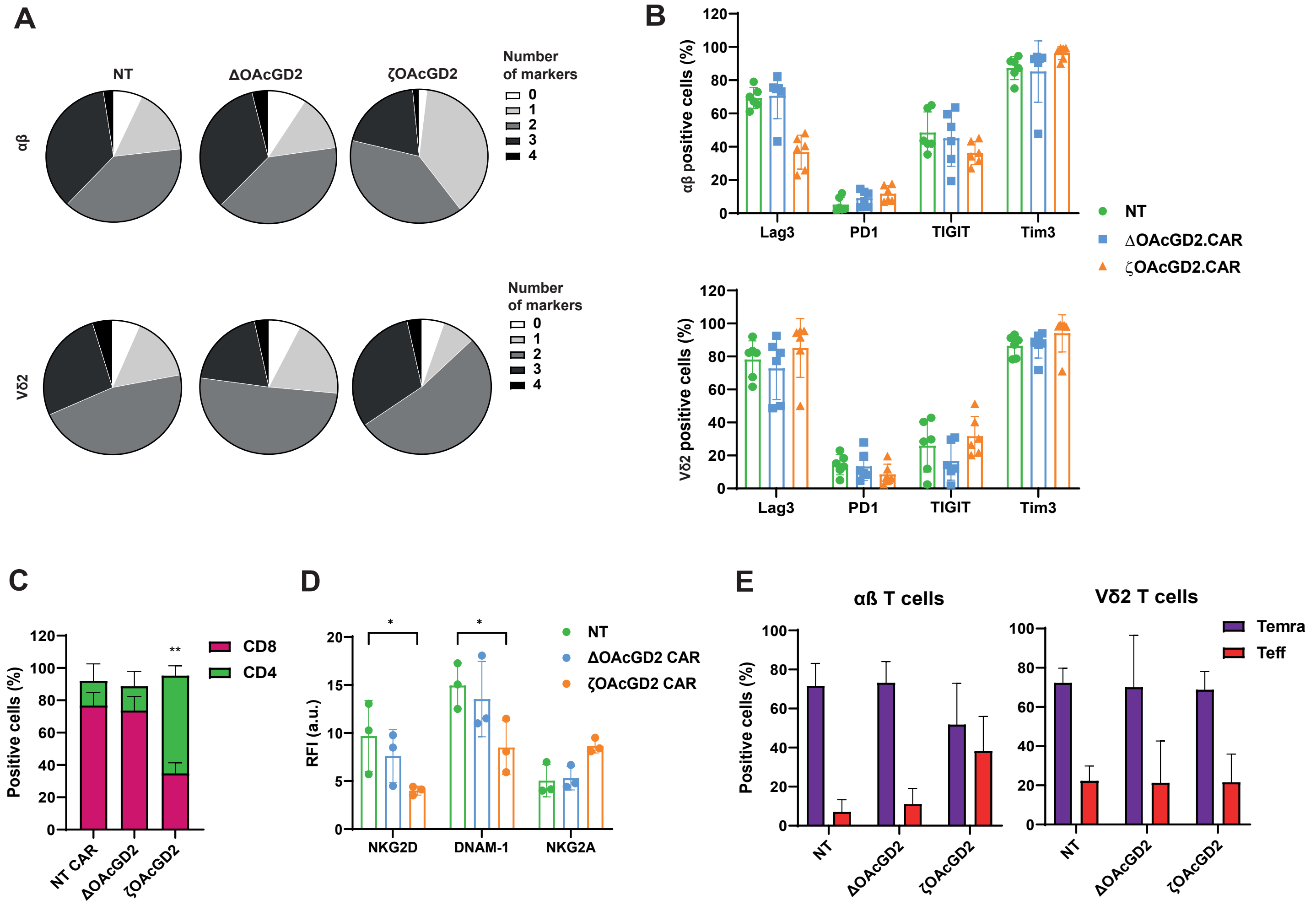


**Extended Figure 1: T cell phenotype is slightly altered by *O*AcGD2-CAR transduction**
**A-B**. Exhaustion profile of αβ (top) and Vδ2 (bottom) CAR-T cells following CAR transduction. Distribution of immune effectors expressing zero, one, two, three or four exhaustion markers (Lag3, PD1, TIGIT and Tim3) (A) and frequency of positive cells expressing exhaustion markers (B) in non-transduced, Δ*O*AcGD2-CAR and ζ*O*AcGD2-CAR αβ T cells (top) or Vδ2 T cells (bottom). Results are presented as % of positive cells, n = 3. **C-E**. Phenotypic profiling of αβ and Vδ2 CAR-T cells following CAR transduction. (C) Expression of CD8 and CD4 were analyzed on non-transduced, Δ*O*AcGD2-CAR and ζ*O*AcGD2-CAR αβ T cells. Results are expressed as % of positive cells, n=3. Two-way ANOVA test, Tukey's multiple comparisons: **, p<0.01. (D) Expression of activating NK-receptors (NKG2D, DNAM-1) and inhibiting NK-receptors (NKG2A) on non-transduced, Δ*O*AcGD2-CAR and ζ*O*AcGD2-CAR Vδ2 T cells. Results are expressed as Ratio of Fluorescence Intensity (RFI) according to isotype, n = 3. Two-way ANOVA test, Tukey's multiple comparisons: *, p<0.05. (E) Memory phenotype of αβ (left) and Vδ2 (right) T cells, in non-transduced, Δ*O*AcGD2-CAR and ζ*O*AcGD2-CAR T cells. Results are presented as % of positive cells, n=3.

### Extended Figure 2: CHLA-01-MED medulloblastoma cell line does not express OAcGD2

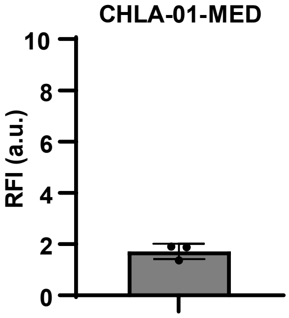


**Extended Figure 2: CHLA-01-MED medulloblastoma cell line does not express *O*AcGD2**
Ratio of Fluorescence Intensity (RFI) of *O*AcGD2 expression in CHLA-01-MED cells. RFI corresponds to the ratio of anti-*O*AcGD2 mAb fluorescence normalized to isotype fluorescence. Results are presented as mean ± SD, n = 3.

### Extended Figure 3: Vδ2 CAR T cells do not display allogeneic reactivity against DIPG-1 in 3D model in contrast to αβ CAR T cells

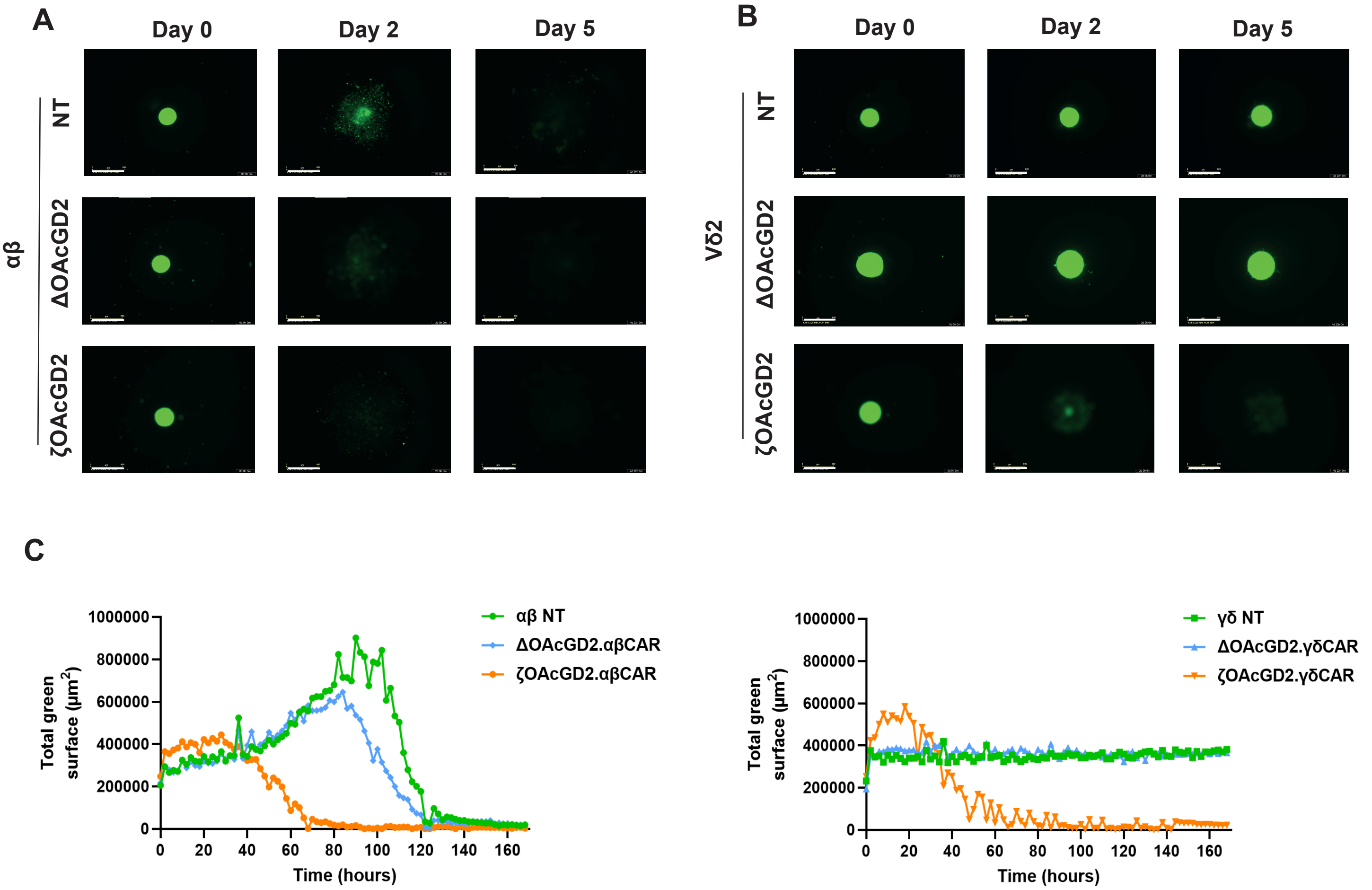


**Extended Figure 3: Vδ2 CAR T cells do not display allogeneic reactivity against DIPG-1 in 3D model in contrast to αβ CAR T cells**

**A-B**. Representative images of tumor cell killing over time against GFP-expressing DIPG-1 cells in 3D-model. Tumor cell killing was monitored over time following addition of αβ (A) and Vδ2 (B) CAR T cells, without transduction or after transduction with Δ*O*AcGD2-CAR and ζ*O*AcGD2-CAR at an effector:target ratio of 3:1. Scale bar = 800 μm **C**. Mean quantification of tumor cell killing over time in presence of non-transduced, Δ*O*AcGD2-CAR and ζ*O*AcGD2-CAR αβ or Vδ2 T cells at an effector:target ratio of 3:1. Results presented represent one representative experiment of 2 independent experiments.
