## Extended Table 1 for "Targeting pediatric High-Grade Gliomas with *O*AcGD2-CAR Vδ2 T cells"

| *Name* | *Cell line/ Primary cells* | *Disease* | *Age* | *Treatment* | *Phase of therapy* | *Disease stage* | *Primary tumor site* | *H3.3K27M status* |
|---|---|---|---|---|---|---|---|---|
| **DIPG-2 (PBT-29)** | Primary cells | DIPG | 12 yrs | NA | No prior treatment | 4 | Pons | Mutant |
| **DIPG-1 (PBT-22)** | Primary cells | DIPG | 5 yrs | NA | No prior treatment | 4 | Pons | Mutant |
| **BT69-NS** | Primary cells | HGG | 14 yrs | 1st line therapy: Stupp* protocol; 2nd line therapy: RAPIRI** protocol | Relapse | 4 | Thalamic | Mutant |
| **BT68 NS** | Primary cells | DIPG | 9 years | NA | Diagnostic | - | Pons | Mutant |
| **BT-35** | Primary cells | HGG | 18 years | 1st line therapy: Stupp* protocol; 2nd line therapy: RT + Irinotecan/ Bevacizumab | Relapse | 4 | Thalamic | - |
| **CHLA-200** | Cell line (adherent) | HGG | 12 years | Chemotherapy and radiation | Post-chemotherapy | 4 | - | - |
| **CHLA-01-MED** | Cell line (neurospheres) | Medullo-blastoma | 8 years | NA | - | - | Brain | - |

**Extended Table 1: Characteristics of pediatric High-Grade Glioma primary cells and cell lines**

*Stupp protocol: Temozolomide chemotherapy and radiation

** RAPIRI: Irinotecan and Rapamycin protocol
