## Supplementary material for "Targeting pediatric High-Grade Gliomas with *O*AcGD2-CAR Vδ2 T cells": STAR Methods

**Resource availability**

Lead contact

Further information and requests for resources and reagents should be directed to and will be

Material availability

| **Reagent or resource** | **Source** | **Identifier** |
| --- | --- | --- |
| **Antibodies** | | |
| Alexa Fluor 647-conjugated F(ab’)2 fragment goat anti-mouse IgG F(ab’)2 fragment | Jackson ImmunoResearch | 115-606-072 |
| anti-human CD107a | BioLegend | 328612 |
| anti-human pan-ab TCR | Beckman Coulter | A39499 |
| anti-human Vδ2 TCR | BioLegend | 331406 |
| anti-human ULBP2,5,6 | BD Biosciences | 748130 |
| anti-human MICA/B | BioLegend | 320907 |
| anti-human HLA-E | BioLegend | 342607 |
| anti-human PVR | BioLegend | 337627 |
| Mouse IgG2a anti-*O*AcGD2 mAb 8B6 | Cerato *et al.* (8) | NA |
| anti-human PD-1 | Miltenyi | 130-120-385 |
| anti-human TIGIT | Miltenyi | 130-116-815 |
| anti-human Lag3 | Miltenyi | 130-120-470 |
| anti-human Tim3 | BD Horizon | 565562 |
| anti-human CD45RA | BD Pharmigen | 550855 |
| anti-human CCR7 | BD Horizon | 566743 |
| anti-human CD62L | BD Horizon | 565040 |
| anti-human NKG2D | BioLegend | 320819 |
| anti-human DNAM1 | BioLegend | 338331 |
| anti-human NKG2A | BioLegend | 375113 |
| anti-human CD8 | BD Pharmigen | 555366 |
| anti-human CD4 | BD Pharmigen | 555349 |
| **Chemicals, peptides, and recombinant proteins** | | |
| EGF | Miltenyi Biotec | 130-097-751 |
| bFGF | Miltenyi Biotec | 130-093-843 |
| N-2 supplement | Life Technologies | 17502-048 |
| B-27 supplement | Life Technologies | 17504-044 |
| Heparin | Sigma-Aldrich | H3149 |
| Peniciline-streptomycin | Gibco | 15140-122 |
| Glutamine | Gibco | 25030-024 |
| Fetal calf serum | Eurobio | CVFSVF00-01 |
| GlutaMAX | Gibco | 35050061 |
| DMSO | WAK-Chemie Medical GmbH | WAK-DMSO-70 |
| *Dynabeads* Human T-Activator *CD3*/*CD28 beads* | Invitrogen | 11131D |
| Zoledronate, Zometa | MedPha Santé, ALTAN | Lot AZ211003 |
| leukoagglutinin (PHA-L) | Sigma-Aldrich | L4144 |
| IL-2 Proleukin® | Clinigen | Lot M000898 |
| Polybrene | Sigma-Aldrich | TR-1003 |
| Monensin | Sigma-Aldrich | M5273 |
| Chromium-51 radionuclide | PerkinElmer | NEZ030002MC |
| Accutase® solution | Sigma-Aldrich | A6964-100ML |
| n-EDTA (0.05%) | Gibco | 25300054 |
| **Critical commercial assays** | | |
| Calcium Phosphate transfection kit | Invitrogen | K278001 |
| Maxima First Strand cDNA Synthesis Kit for RT-qPCR | Thermo Scientific | K1642 |
| **Experimental models: cell lines** | | |
| CHLA-200 | Children Oncology Group, Philadelphia | NA |
| CHLA-01-MED | ATCC | CRL-3021 |
| PBT-22 (DIPG-1) | Fred Hutchinson Cancer Research Center, Seattle | NA |
| PBT-29 (DIPG-2) | Fred Hutchinson Cancer Research Center, Seattle | NA |
| BT35 | LBP of Strasbourg, France | NA |
| BT68NS | LBP of Strasbourg, France | (13) |
| BT69NS | LBP of Strasbourg, France | (13) |
| **Softwares** | | |
| Prism 9.0 GraphPad | Graphpad Software | https://www.graphpad.com/ |
| FlowJo V10 | FlowJo | https://www.flowjo.com/ |
| IncuCyte® S3 Analysis software | Sartorius | https://www.sartorius.com/en |
| **Others** | | |
| DMEM high glucose media | Gibco | 21969 |
| DMEM F-12 media | Gibco | 21331-020 |
| Lymphocytes separation medium | Eurobio | CMSMSL01-0U |
| Hitrap rProtein A FF column | GE Healthcare Bios-Sciences, Uppsala, Sweden | 17507901 |
| Ultra Low Adhrence Plate (ULA) | Corning | 7007 |

**Experimental Model and Subject Details**

Tumor cells and irradiation

The human DHG CHLA-200 cell line was obtained from the Children's Oncology Group Cell Culture and Xenograft Repository (Philadelphia, PA, USA). Human primary DIPG PBT-22 called DIPG-1 and PBT-29 called DIPG-2 were obtained from the Fred Hutchinson Cancer Research Center (Seattle, WA, USA). Human primary DIPG cells BT68NS, BT69NS and BT35 were kindly provided by Natacha Entz-Werlé at LBP of Strasbourg, France (12). The human medulloblastoma cell line CHLA-01-MED was obtained from ATCC (Manassas, VA, USA). CHLA-200 cell line was grown in DMEM with 10% heat-inactivated fetal calf serum, 2 mM L-Glutamine, 100 units/mL penicillin, and 100 µg/mL streptomycin, at 37°C in 5% CO_2_. PBT-22, PBT-29 and CHLA-01-MED were grown in Neurobasal medium [DMEM/Ham F12, 2 mM L-glutamine, N2- and B27-supplements, 2 mg/mL heparin, 20 ng/mL EGF and 25 ng/mL bFGF, and 100 IU/mL penicillin and 100 mg/mL streptomycin] at 37°C in 5% CO_2_. BT68NS and BT69NS cells were grown in DMEM/Ham F12, B27-supplement, 2 mM GlutaMAX, 20 ng/mL EGF and 20 ng/mL bFGF. BT35 primary cell was grown in DMEM/Ham F12 GlutaMAX-supplemented medium with 10% heat-inactivated fetal calf serum. Cells were irradiated at a rate of 1.6 Gy/minute by a Faxitron cabinet X-ray system model CP160. All experiments were performed with cells in culture for less than 3 months or passage < 20, and cells were regularly checked for mycoplasma contamination.

T cell isolation and expansion

Peripheral blood mononuclear cells (PBMCs) were obtained from seven blood donors (DO240, DO3791, D3.1, D7314, D2519 and DO5558) at the Etablissement Français du Sang (EFS) with informed consent (Blood product transfer agreement relating to biomedical research protocol 97/5-B—DAF 03/4868). PBMCs from donors were isolated using density gradient centrifugation with Lymphocytes separation medium, then frozen in human SVF/10% DMSO and stored in liquid nitrogen. For αβ T cell selection, PBMCs were stimulated with *Dynabeads* Human T-Activator *CD3*/*CD28 beads*. For Vδ2 T cell expansion, PBMCs (1.5.10^6^/ml) were stimulated with 5µM Zoledronate in 24-well plates and cultured at 37°C in 5% CO_2_. At day 3, cell cultures were supplemented with 100 IU IL2/ml. At day 14, Vδ2 T cell cultures were examined by flow cytometry for γ9δ2 T cell expansion and purity. Vδ2 T cells represented at least 90% of the cells in the culture.

**Method Details**

Retroviral vector production and T-cell transduction

Plasmids were obtained by GeneCust (Luxembourg). Retroviral supernatants were produced by transfection of Phoenix-Ampho packaging cells. Two million Phoenix-Ampho cells were seeded in 10-cm-diameter dishes 24 h prior to transfection. Transfection was performed with 15 µg pMXA/O-Acetyl-GD2 CAR plasmid DNA using Calcium Phosphate transfection. The medium (10 ml) was replaced 6 h after transfection. The conditioned medium was collected 48 h post-transfection, filtered through 0.45-µm pore-size filters and kept at -80°C until use. For primary αβ T cells, PBMC were stimulated with *Dynabeads* Human T-Activator *CD3*/*CD28 beads* and transduced four days after stimulation. For selected Vδ2 T, cultures were stimulated with irradiated (35 Gy) pooled allogenic feeder cells, 1 µg/ml leukoagglutinin (PHA-L), and IL-2 (300 IU/ml). Four days after stimulation, T cells were resuspended in RPMI 1640 culture medium supplemented with 8% pooled allogeneic human serum and 100 IU/mL of recombinant IL-2, seeded at 1 × 10^6^ cells in 1 mL per well into 6-well plates, and exposed to 2 × 2 mL of retroviral supernatant by spinoculation (2400 g, 1.5 h, and 32°C) in the presence of 4 μg/mL polybrene. The culture medium was changed 24 h after infection. Mock (non-transduced) controls were performed in parallel, by which the supernatant of untransfected packaging cells was added to primary T cells. The transduction efficiencies were assessed 5 days later by staining the CAR with a Alexa Fluor 647-conjugated F(ab’)2 fragment goat anti-mouse IgG F(ab’)2 fragment. By 20 days after transduction, T cells were sorted by flow cytometry on TCR and CAR expression.

RT-qPCR

RNAs were extracted from cell pellets using Nucleospin RNA Plus kit (Macherey Nagel, Düren, Germany) according to the recommendations. The quantity and quality of RNA were respectively evaluated using the NanoDrop® ND-1000 spectrophotometer (Nanodrop Technologies, Wilmington,DE) and the Agilent 2100 Bioanalyser (Agilent, SantaClara, CA). RNA integrity number (RIN) was > 7.5 in all cases.

RNA reverse transcription was performed using Maxima First Strand cDNA Synthesis Kit for RT-qPCR (Thermo Scientific, MA). Quantitative real-time PCR (qPCR) assays were performed in duplicates using the Perfecta™ SYBR® Green FastMix™, Low ROX™ (Quanta BioSciences, CA) and the real-time thermal cycler qTower3 (Analytik Jena AG, Germany). A melting curve was performed at the end of each qPCR. Linearity and efficiency of the qPCR were checked for each gene with a standard curve of 4 logs. Efficiency was >90% in all cases. Fold change was calculated with Pfaffl MW formulae using a RNA extracted from a glioblastoma cell line as calibrator and three housekeeping genes RPLPO, TATA, and GAPDH (n = 4).

Table 1: Sequences of Forward and Reverse primers

| Gene | Forward | Reverse |
| --- | --- | --- |
| MICA | CTGGGAAATAAGACATGGGACAG | GAATGCAAGCCTTCTTTCTGGT |
| MICB | CAGAGACCGAGGACTTGACAG | GAATGCAAGCCTCCTTTCTGGT |
| BTN2A1 | GTCATACACAACATCACAGCCC | CTTAGAGCCTAGTCCTGCCAC |
| BTN3A1 | GAATACACAACGTCACAGCCT | ATCCCTCCATCCTTGTAACCC |
| ULBP2 | GCCGCTACCAAGATCCTTCTGTG | CTTGAACCGCACACCACCGT |
| RPLPO | GATTACACCTTCCCACTTGCT | TAGTCAAAGAGACCAAATCCCA |
| GAPDH | GAAGGTGAAGGTCGGAGTC | GAAGATGGTGATGGGATTTC |
| TATA | CAAGAGTGAAGAACAGTCCAG | ACAAGGCCTTCTAACCTTATAGG |

CD107a cell surface mobilization

Tumor cells were cocultured with T cells at 1:1 effector:target ratio in defined culture medium containing 5 μmol/L monensin and APC-labeled anti-human CD107a mAb for 4 hours at 37°C in a humidified atmosphere with 5% CO_2_. For CD107a assay with irradiated cells, coculture was made with conditioned media. αβ and Vδ2 T cells were then stained with PE-labeled anti-human pan-αβ TCR mAb and FITC-labeled anti-human Vδ2 TCR mAb respectively and analyzed by flow cytometry. Acquisition was performed using Accuri C6Plus flow cytometer (BD Biosciences), and the events were analyzed using the FlowJo software 10 (BD Biosciences).

Cell surface phenotyping

Tumor cell surface phenotype was evaluated by flow cytometry using anti-ULBP2,5,6 ; anti-MICA/B ; anti-HLA-E and anti-PVR. Mouse IgG2a anti-*O*AcGD2 mAb 8B6 was obtained as described previously (8,21) and purified using Hitrap rProtein A FF column. *O*AcGD2 expression on tumor cell is measured with 8B6 clone at 10 µL/mL on dissociated tumor cells (by 1X accutase for neurospheres or 1X trypsin for spheroids). T cell surface phenotype was evaluated by flow cytometry using anti-PD-1 ; anti-TIGIT ; anti-Lag3 ; anti-Tim3 ; anti-CD45RA; anti-CCR7 ; anti-CD62L ; anti-NKG2D ; anti-DNAM1 ; anti-NKG2A ; anti-CD8 and anti-CD4. Acquisition was performed using Accuri C6Plus flow cytometer (BD Biosciences), BD FACSCanto™ II Clinical Flow Cytometry System and BD FACSymphony™ A5 Cell Analyzer and the events were analyzed using the FlowJo software 10 (BD Biosciences).

Cytolytic activity

Cytolytic activity was assessed through standard ^51^Cr-release assay. pHGG cultures were labeled with ^51^Cr (75 μCi for 1 × 10^6^cells) for 1 hour at 37°C, washed three times with define culture medium, plated at 3 × 10^3^ cells per well, and T cells were added at different effector:target ratios in 96-well round-bottom plates. After a 4-hour coculture at 37°C, tumor cell lysis was measured in supernatants using a scintillation counter. Percentage of tumor target cell lysis = [(experimental release-spontaneous release) / (maximum release-spontaneous release)] × 100. Maximum and spontaneous releases were determined by adding 1% Triton X-100 (Sigma) or medium, respectively.

Videomicroscopy

The IncuCyte® Live-Cell Imaging System was used to monitor spheroid surface. Cells were seeded at 10,000 cells/well in Ultra-Low Adherence 96-wells plate and allowed to grow for 48 hours. After spheroid formation, 100 µL of medium were removed and T cells were added at 3:1 effector:target ratio with 300 IU/mL of IL-2. Spheroid surface was analyzed by the IncuCyte® S3 Analysis software.

**Statistical analysis**

Results are presented as mean ± SD of three or four independent experiments, unless otherwise noted. For statistical analyses, Student t-test or two-way ANOVA were performed using Prism 7.0 GraphPad Software. P-value below 0.05 was considered significant: * p<0.05, ** p<0.01, *** p<0,001, **** p<0.0001. Sample sizes of independent experiments (n) are reported in the corresponding figure legends.
